# The *NDE1* genomic locus can affect treatment of psychiatric illness through gene expression changes related to MicroRNA-484

**DOI:** 10.1101/087007

**Authors:** Nicholas J. Bradshaw, Liisa Ukkola-Vuoti, Maiju Pankakoski, Amanda B. Zheutlin, Alfredo Ortega-Alonso, Minna Torniainen-Holm, Vishal Sinha, Sebastian Therman, Tiina Paunio, Jaana Suvisaari, Jouko Lönnqvist, Tyrone D. Cannon, Jari Haukka, William Hennah

## Abstract

Genetic studies of familial schizophrenia in Finland have observed significant associations with a group of biologically related genes, *DISC1, NDE1, NDEL1*, *PDE4B* and *PDE4D,* the “DISC1 network”. Here, we utilize gene expression and psychoactive medication use data to study their biological consequences and potential treatment implications. Gene expression levels were determined in 64 individuals from 18 families, whilst prescription medication information has been collected over a ten-year period for 931 affected individuals. We demonstrate that the *NDE1* SNP rs2242549 associates with significant changes in gene expression for 2,908 probes (2,542 genes), of which 794 probes (719 genes) were replicable. A significant number of the genes altered were predicted targets of microRNA-484 (p=3.0×10^−8^), located on a non-coding exon of *NDE1.* Variants within the *NDE1* locus also displayed significant genotype by gender interaction to early cessation of psychoactive medications metabolized by CYP2C19. Furthermore, we demonstrate that miR-484 can affect the expression of *CYP2C19* in a cell culture system. Thus, variation at the *NDE1* locus may alter risk of mental illness, in part through modification of miR-484, and such modification alters treatment response to specific psychoactive medications, leading to the potential for use of this locus in targeting treatment.

## Introduction

The identification of genes that predispose to complex psychiatric traits is an important aspect in studying these conditions, however it is vital that this information is then used to improve our biological understanding and ultimately the treatment procedures for the disorders. This can be achieved through genetic studies in which, instead of using an end state diagnosis, alternative traits are employed that can measure a biological or pharmacological aspect of the condition.

Polygenic disorders, such as schizophrenia, are influenced by numerous interacting genetic factors, therefore identification of one candidate gene may aid in identification of others. This approach has been used in a large Finnish family cohort, in which *DISC1* (Disrupted in Schizophrenia 1) was previously associated with schizophrenia (1, 2), and which led to observation of association with four other genes *(NDE1, NDEL1, PDE4B,* and *PDE4D)* (3, 4) that encode protein binding partners of the DISC1 protein (5-8). The idea that such protein interaction partners of DISC1 are encoded for by genes which show genetic interaction in mental illness is termed the DISC1 network hypothesis. Specifically, multiple associations for psychiatric (2, 9-11) and related endophenotypes, including memory (12), cognitive, and neuroimaging (13) phenotypes, have been reported for *DISC1* in Finnish cohorts. By conditioning genome-wide linkage data for schizophrenia on *DISC1,* a peak of linkage at chromosome 16p was observed (3), near to *NDE1* (Nuclear Distribution Element 1). This was followed up through association analysis at the *NDE1* locus, leading to the observation that a haplotype and its constituent SNPs associate with schizophrenia in this cohort, in a gender dependent manner (3). Genetic association for schizophrenia was therefore tested for other DISC1 binding partners in this family cohort (4). Although SNPs and haplotypes from six other genes were initially observed to associate, only variants in *NDEL1* (NDE-like 1, a close paralogue of *NDE1)* and in the phosphodiesterases *PDE4B* and *PDE4D* replicated when tested in a second, distinct sample from the cohort (4). Recently, through further investigation of the roles of these variants in the DISC1 network, the *NDE1* locus has been identified to increase risk to schizophrenia in this Finnish family cohort through interaction with high birth weight, a promising proxy measure for multiple pre- and/or perinatal environments (14).

The role of the DISC1 network as a source for genetic risk for neuropsychiatric disorders is controversial due to the absence, to date, of evidence for their involvement in population based genomic studies of common variation (15, 16). However, these genes have been implicated at least within specific populations through strong evidence emerging from family based approaches and the studies of rare variants. In addition to the evidence from the Finnish family cohort, the *DISC1* and *PDE4B* genes are disrupted by chromosomal aberrations in Scottish families with major mental illness (8, 17, 18). Furthermore, *NDE1* is independently implicated in major mental illness through its presence at 16p13.11, which is subject to duplications in schizophrenia (19-22), as well as being directly implicated through rare SNPs in patients (23). The importance of the NDE1 protein for neurodevelopment more generally has been dramatically demonstrated in individuals with biallelic loss of the functional *NDE1* gene, leading to severe microcephaly phenotypes, sometimes described in conjunction with lissencephaly or hydrocephaly (24-27). Deletion of only one copy of the 16p13.11 locus, meanwhile, has been associated with neurological conditions including autism and epilepsy (28). Recently it has been shown that expression of mature miR-484, a microRNA that is encoded on an untranslated exon of *NDE1,* led to alterations in neural progenitor proliferation and differentiation, as well as behavioural changes in mice, thus implicating the microRNA in the phenotypes associated with 16p13.11 duplication (29).

We have previously studied the effect of DISC1 network genetic variation on gene expression in a publicly available population cohort of the CEU (Utah residents with North and Western European ancestry) individuals, with 528 genes being differentially expressed across 24 variants studied, of which 35 genes had pre-existing supporting evidence for a role in psychosis (30). Intriguingly, seven of these affected genes were noted to be targets for drugs prescribed for psychiatric illness, leading to the hypothesis that these DISC1 network variants, through their action on gene expression, may alter treatment outcome for medications designed to target these genes (30).

Here, in order to advance our understanding of the role these genes play in the aetiology of schizophrenia in Finland, we take this approach further. This is accomplished by utilizing data on gene expression levels in case families in which these DISC1 network genetic variants have been previously demonstrated to associate with schizophrenia (1-4), as well as by using data collected on how different psychoactive medications are used by the affected individuals within these families.

## Methods

### Study Samples

The principal samples used here are part of a larger study of familial schizophrenia. These are Finnish patients born between 1940 and 1976, who were identified through the hospital discharge, disability pension, and the free medication registers (1, 31). The cohort totals 458 families (498 nuclear families) that contain 2,756 individuals, of whom 2,059 have been previously genotyped for the DISC1 network genes (1-4). Of these genotyped individuals, 931 are classified as affected with major mental illnesses using criteria from the Diagnostic and Statistical Manual of Mental Disorders, fourth edition (DSM-IV) (32). These include 635 diagnosed with schizophrenia, 125 with schizoaffective disorder, 95 with schizophrenia spectrum diagnoses, and 76 with other mental illness including bipolar disorder and major depression. Here, two sub-sets of this familial sample were used as a discovery cohort (18 families, N=64) to study gene expression level changes, and all affected individuals were used as a discovery cohort (N=931) to study medication use.

In order to replicate the gene expression results obtained from this family data, two independent cohorts were used. The first of these replication cohorts was a Finnish discordant twin pair sample ascertained for schizophrenia (N=73), for which information about recruitment and clinical evaluation has been described previously (33). Briefly, the participants are 18 schizophrenia patients, their 18 unaffected co-twins, and 37 control twins who have provided blood samples for gene expression analysis (N=73). The second replication cohort was the Genotype-Tissue Expression (GTEx) database (N=338), a publicly available resource for exploring the correlation between genotypes and gene expression across multiple tissues and in a genome-wide manner (accessed on September 2^nd^ 2016, www.gtexportal.org/home) (34). To best match the source of the RNA used in the discovery cohort studies, data from whole blood was used for the GTEx tests.

### Gene Expression Data

Total RNA was extracted from fresh blood samples from 82 individuals, with 18 individuals excluded from further analysis as their samples RNA Integrity Number (RIN) were lower than 8. These individuals are from 18 families that were re-approached to provide RNA for gene expression analysis based on prior genetic observations in these families including *DISC1* (1), *RELN* (35) and *TOP3B* (36). Genome-wide gene expression measures were assayed for this discovery cohort using Illumina HumanHT-12 v4.0 Expression BeadChip. Of the 48,212 probes on the chip, 11,976 were significantly detectable at a threshold of p≤0.01 in more than 90% of individuals. The expression data for these probes were processed using quantile normalization followed by log_2_ transformation. Raw anonymous data regarding this family cohort can be accessed at the Gene Expression Omnibus (GEO) database (GSE48072). For the replication twin cohort (N=73), genome-wide gene expression data has been measured using Illumina Human WG6 v3.0 chip, as reported previously in detail (37). After quality control and data processing, identical to that used on the family data, 18,559 probes from this chip were significantly detectable.

### Genotyping

In the discovery sub-cohorts used here, both genotype and expression data was available from 39 individuals, while 931 individuals had both genotype and medication data available. Thus, in order to ensure sufficient numbers of individuals for statistical testing, we only studied genetic variants that met specific minor allele homozygote frequencies in these sub-cohorts. In the discovery cohort for gene expression, a cut-off value for the minor allele homozygote frequency of ≥10% was implemented, providing five variants *(DISC1:* HEP3 haplotype [comprising SNPs rs751229 and rs3738401] and rs821616; *NDE1:* rs4781678, rs2242549, and rs1050162) with which to perform the analysis. For the discovery cohort for medication use, the frequency of the genetic variants was restricted to those with a minor allele homozygote frequency of at least 5%. This allowed seven variants from three DISC1 network genes to be studied *(DISC1:* rs821616; *NDE1:* rs4781678, rs2242549, rs881803, rs2075512, and the haplotype of the four SNPs *‘NDE1* Tag haplotype’; *PDE4B:* rs7412571)

The genotypes for the replication cohort of discordant twins were produced with the same method and at the same time as those described previously (1, 3, 4), with only two variants analysed *(NDE1:* rs2242549 and rs1050162) using the gene expression data. The analysis using the GTEx database as a replication cohort was conducted for all variants studied in the families, except for the *DISC1* haplotype.

All genetic variants analysed in this study have been implicated by previously described evidence as being associated with the aetiology of schizophrenia in this cohort (1-4), with variants in both *DISC1* and *NDE1* having displayed prior gender dependent effects (1, 3, 12). Therefore, no multiple test correction has been applied to correct for the multiple testing across variants or gender interaction models, as they can all be considered hypothesis based. However, since we are screening alternative phenotypes in a hypothesis free manner, we have applied the measures described in the following sections in order to further characterise these *a priori* variants.

### Association between Genome-Wide Expression Levels and Genotypes

For the discovery cohort used to study gene expression, a mixed effect linear regression model was fitted for each probe and genotype using R (RStudio version 0.99.489) lme4 package (38), after correcting for gender, age, affection status, and family or twin status effects as a covariate. This analysis was performed separately but identically for the discovery cohort and the replication twin cohort. We used the false discovery rate (FDR) method in place of a family-wise error rate (FWER). FDR is widely applied for microarray analyses because it allows more genes to be extracted for further exploration, and were performed using the qvalue package in R (39) to estimate the FDRs of q≤0.05. The post hoc power of our small familial discovery cohort to detect gene expression changes was estimated using R package ssize.fdr (40). The GTEx database was mined using its own in-built test procedure, entering in a list of gene IDs to be tested against our SNPs of interest. Data from whole blood was used in order to replicate only those genes identified as significantly altered in their gene expression levels at a cut-off of p≤0.05. When testing for replication probes significant (p≤0.05) in the discovery cohort prior to application of FDR were studied. See Figure 1 for a flow chart of analysis.

**Figure 1:**
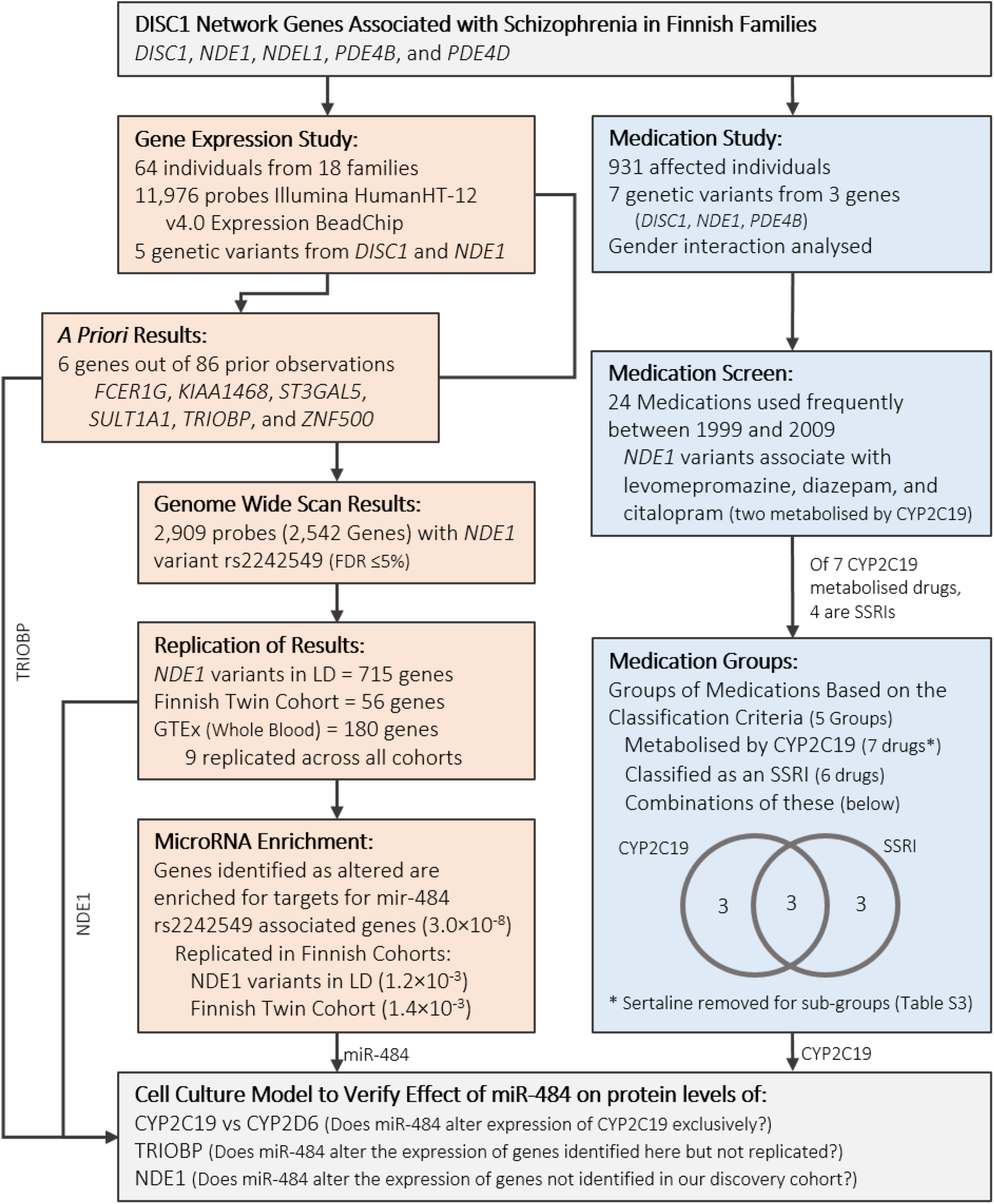
Schematic flow chart of the analysis undertaken in this study.

### MicroRNA Target Prediction and Enrichment Analysis

A comprehensive list of predicted 3’UTR targets for miR-484 were obtained from the miRWalk database (41), considering only genes predicted by at least six of the 12 programs. Ingenuity Pathway Analysis (QIAGEN Redwood City, www.qiagen.com/ingenuity) was used to analyse potential enrichment of these genes amongst those significantly altered in their expression levels by *NDE1* SNPs in the discovery and replication cohorts.

### Medication Data

Prescription medication data from the 931 affected individuals of the schizophrenia family cohort for the period 1^st^ January 1996 to 31^st^ December 2005 was obtained from the Finnish National Prescription Register of the National Social Insurance Institution (SII) (42), thus forming the discovery cohort for medication use. In this cohort “medication use” is based on purchases of prescribed psychoactive drugs for which the SII have paid a reimbursement. Data from this register contains information on date of purchase and the dose, stated as the international standard daily defined dose. Thus using this data medication periods were defined according to method 4 proposed by Mantel-Teeuwisse et al, multiplying defined daily dose by a factor of 1.1 and filling 15-day gaps between medication periods (43). This medication period data was used to determine the probability of cessation of each drug by genotype and converted into a binary variable using three months as a cut-off. This three months cut-off reflects that an individual either purchased more of the same medication, or purchased a different medication within a three month period after the original purchase date. Only psychoactive medications with at least 15 instances of use for three months or less were taken for association analysis of the individual drugs, meaning that, of the possible 101 psychoactive drugs for which data was available, only 24 were analysed. When data from multiple drugs were combined in order to study classes of medication, all medications for which data is available were used, regardless of individual frequency. In order to account for the fact that some medication periods may come from the same individual, analysis of the medication usage used logistic regression with GEE-estimation (Generalized Estimating Equations), as utilized by the geepack-package for R (44, 45). Bonferroni correction was used to correct for the multiple tests in the analysis of the individual medications (24 tests) and the groups of medications (five tests). While p-values are presented unadjusted, only those that would surpass the Bonferroni correction based thresholds (p≤0.0021 and p≤0.01 respectively) are highlighted. See Figure 1 for a flow chart of analysis.

### Cell Culture and Western Blotting

To determine whether miR-484 had the potential to affect the expression of selected proteins in a human cell based system, NLF neuroblastoma cells (Children’s Hospital of Philadelphia) were grown in RPMI 1640 / 10% foetal calf serum / 2mM L-Glutamine (all from Thermo Fischer Scientific) and transfected with 50nM of either a mimic of mature miR-484 (QIAgen, Sy-hsa-miR-484) or a negative control microRNA (QIAgen, AllStars Negative Control microRNA) using Lipofectamine 2000 (Thermo Fischer Scientific) according to manufacturer’s instructions. After 48 hours, cells were lysed using PBS / 1% Triton X-100 / 20mM MgCl_2_ containing protease inhibitor cocktails and DNaseI. Lysates were Western blotted and proteins detected using the following antibodies: anti-α-actin (Sigma, A2066), anti-CYP2C19 (Novus Biologicals, NBP1-19698), anti-CYP2D6 (Abnova, H00001565-B01P), anti-NDE1 (ProteinTech, 10233-1-AP), anti-TRIOBP (Sigma, HPA019769) and anti-α-tubulin (Sigma, T9026). IRDye secondary antibodies were used (LI-COR) and the signal visualized and quantified using an Odyssey CLx infrared imaging system (LI-COR) and associated software. All membranes were probed with secondary antibodies alone first to ensure specificity of signal. Antibody signals were normalized to actin as a loading control. Mean fold-changes between the control and miR-484-treated samples were calculated from 7-8 internal replicates. Three independent experiments were performed and the results compared by two-tailed paired Student’s t-test.

## Results

### Replication of Gene Expression Changes from Previous Studies

In our previous analysis of gene expression in association with DISC1 network variants using publicly available data on the CEU (Utah residents with North and Western European ancestry) individuals, 86 genes were found to be differentially expressed (30). To verify these results, five of variants were tested again, this time using the Finnish family cohort. In total, six of the gene expression changes previously reported could be replicated, for the genes *TRIOBP, ZNF500, KIAA1468, FCER1G, SULT1A1,* and two probes for the *ST3GAL5* gene. These were all in association with the status of the *NDE1* gene locus (rs2242549).

### *Differentially Expressed Genes Associated with* DISC1 *Pathway Genotype*

To further investigate the effect of DISC1 network variants on gene expression, we used this Finnish family cohort as a discovery sample to investigate the association of previously positive *DISC1* and *NDE1* variants with genome-wide gene expression of 11,976 probes. Notably, the *NDE1* SNP rs2242549 was significantly associated with gene expression levels of a large number of the probes (Tables 1 and S1, Figure S1). Specifically, 3,824 probes representing 3,314 distinct genes showed uncorrected association with the *NDE1* SNP rs2242549 (p≤ 0.05), of which 2,908 probes, representing 2,542 distinct genes were associated at FDR ≤5 %. We also verified that, despite the size of the discovery cohort, it had sufficient power to detect the differentially expressed genes associated with the rs2242549 variant (Figure S2). In contrast, no probes were significantly altered in expression levels in association at FDR ≤5 % with either the *DISC1* variants tested (SNP rs821616 of the “HEP3” haplotype comprising rs751229 and rs3738401) or with the other *NDE1* SNPs tested (rs4781678 or rs1050162).

**Table 1:**
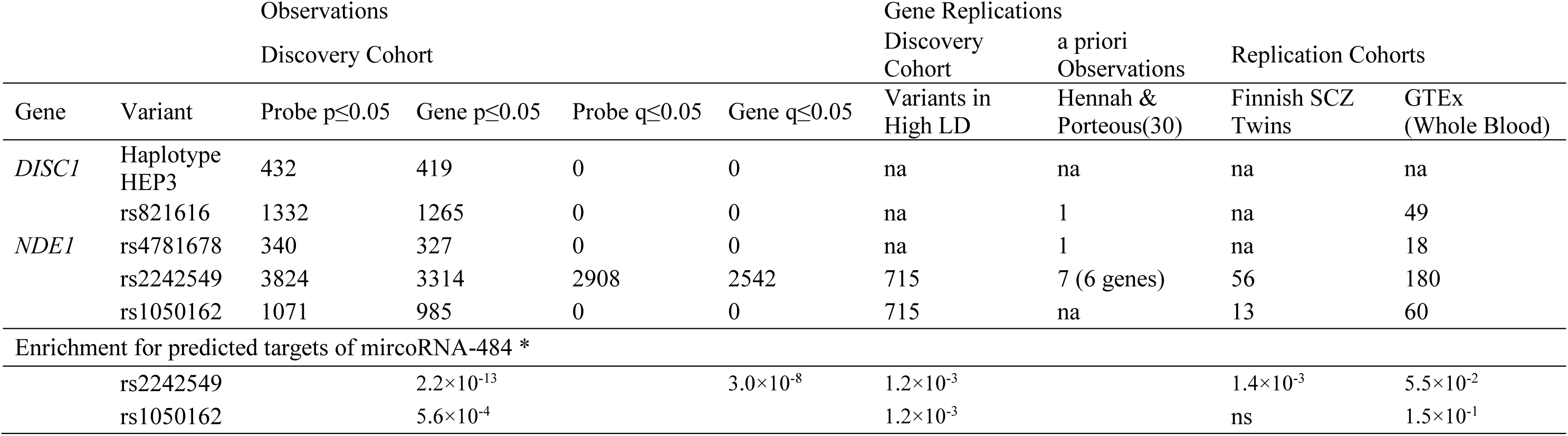
Number of probes and genes significantly altered by variants in the DISC1 network, how they replicate across different cohorts, and how they overlap with the predicted targets of miR-484. ^*^Targets predicted by at least 6 out of 12 prediction tools summarised by miRWalk were uploaded to Ingenuity Pathway Analysis (IPA), thus enabling a test for enrichment when the subsequent probe/gene lists were studied. na = value was not applicable as it was not tested, ns = value was not significant nor returned by IPA.

### *Replication of Gene Expression Changes Associated with* NDE1 *Genotypes*

In order to replicate these observations, we pursued three lines of supporting evidence. Firstly, we noted that the *NDE1* SNPs rs2242549 and rs1050162 were in high linkage disequilibrium (LD) in the full family cohort (r^2^=0.88, n=1891 individuals genotyped at both loci), and therefore can be assumed to act as an internal replication of observations. Thus, within the discovery cohort, of the genes whose expression levels were associated with rs2242549, 752 probes representing 695 genes were also significantly associated with rs1050162 (p≤0.05). Secondly, we utilized existing data from a twin cohort for schizophrenia from Finland (33, 37) as a replication cohort, identifying 56 probes, each representing a different gene, with replicable significant alteration in their expression levels associated with rs2242549 (Tables 1 and S1). Finally, we used the publicly available database GTEx as an additional independent replication cohort, from which we were able to directly test 2,651 of the 3,314 genes identified (those probes with an official gene name), confirming that 180 genes also display significant gene expression changes related to the *NDE1* SNP rs2242549 in this database (Tables 1 and S1).

In total, 794 probes representing 719 genes had supporting evidence from at least one additional source for their changes in expression related to the *NDE1* variant rs2242549, of which 76 probes from 73 genes had supporting evidence from more than one source, and 4 probes from independent genes *(ITGB5, OVGP1, PGRMC1, TST)* had supporting evidence from all three sources, that is SNPs in high LD in the discovery cohort, as well as independent replication cohorts of Finnish twins and the GTEx database.

### *Enrichment of miR-484 Target Genes Amongst Genes Whose Expression is Associated with* NDE1 *Genotype*

The finding that expression levels of such a large number of genes could be altered by a single genetic locus was surprising, especially given that the principal functions of the NDE1 protein are not known to be in gene regulation. The *NDE1* locus also encodes for a microRNA (miR-484), which is located on a non-coding 5’ exon of the longest splice variant of the *NDE1* gene. Since the major function of microRNAs is in the regulation of expression of other genes it is the most likely explanation for the sheer number of expression changes observed to associate with these SNPs. We therefore investigated whether the set of genes whose expression is altered by these *NDE1* SNPs overlapped with those genes predicted to be targets of miR-484.

Using the miRWalk database (41), 16,027 gene targets are predicted for miR-484, of which 2,588 are predicted by at least six or more of the 12 independent prediction programs used in the database (data collated June 2016). Upon examining the list of genes whose expression is altered by SNPs in the *NDE1* locus at FDR ≤5 % in our discovery cohort, these probes were indeed seen to be enriched for predicted targets of miR-484 (p=3×10^−8^). Employing the same tests to the three replication studies described above, the enrichment in miR-484 targets was also present for the set of these genes whose expression level is significantly associated with the genotype of both *NDE1* SNPs rs2242549 and rs1050162 (which were in high LD), and amongst those genes which could also be observed in the replication twin cohort. In contrast, they were not enriched amongst those genes replicated by data from the GTEx database (Table 1).

### *Medication Cessation Associated with* NDE1 *Genotype*

In our previously published analysis we found *NDE1* genotypes that significantly associated with early cessation of particular medications relevant to mental health (30). We therefore tested whether *NDE1* genotypes were also associated with cession of specific medications in a discovery cohort consisting of all affected individuals from the Finnish family cohort. Screening all of the medications frequently used within this discovery cohort, we observed an association between *NDE1* rs4781678 genotype and early cessation of use of the antipsychotic levomepromazine (OR=4.13 per C allele; 95%CI=1.72-9.91; p=0.00090). When analysed in interaction with gender, association was further observed across *NDE1* genotypes with early cessation of the use of diazepam and citalopram (Table S2), two drugs that share a common principal metabolizing enzyme, CYP2C19 (Cytochrome P450 2C19, Table S3) (46-49). We therefore asked whether *NDE1* genotype was associated with the subset of medications metabolized by the CYP2C19 enzyme. Since four out of seven of the drugs metabolized by CYP2C19 were selective serotonin reuptake inhibitors (SSRIs), we studied these as a separate group as well as further separated based on CYP2C19 metabolism. No significant interaction was observed for the grouping of all drugs metabolized by CYP2C19, however, a genotype by gender interaction was noted when all SSRIs were grouped together (rs2075512, OR=0.37; 95%CI=0.17-0.79; p=0.010). When all SSRIs metabolized by CYP2C19 were tested, this genotype by gender interaction became significant (ranging from: OR=0.27 to 0.31; 95%CI=0.11 to 0.13–0.64 to 0.71; p=0.0030 to 0.0060) for four out of the five *NDE1* markers tested, while no interaction was noted for SSRIs not metabolized by CYP2C19 (Table 2 and Figure S3). We analysed the remaining drugs metabolized by CYP2C19 together as another grouping (“non-SSRIs metabolized by CYP2C19”). Interestingly, a significant interaction was observed (ranging from: OR=3.33 to 5.82; 95%CI=1.44 to 2.25–7.25 to 15.0; p=0.0013 to 0.00030) for all five *NDE1* markers (Table 2 and Figure S3). In this case, however, the gender effect is reversed, with SNPs being associated with cessation among females, in contrast to SSRIs metabolized by CYP2C19, for which SNPs were associated with cessation among males.

**Table 2:** Results for the association of *NDE1* variants with groups of medications based on their metabolism by the CYP2C19 enzyme and/or Selective Serotonin Reuptake Inhibitor (SSRI) class status, showing the p-values and odds ratios (and 95% confidence intervals) for the interaction model. P-values ≤0.01 are below the Bonferroni correction threshold for the five groups tested. P-values and their respective ORs that are below the Bonferroni threshold are in bold. Medications included in each group analysis can be found in Figure S3 and Table S3.

| *NDE1* Variant | SNP<br>p-value | SNP*Gender<br>p-value | OR (95% CI) |
| --- | --- | --- | --- |
| All psychoactive drugs metabolised by CYP2C19 (N=510-581, no of instances: 843-945) |  |  |  |
| **rs4781678** | 0.79 | 0.90 | 0.97 (0.55 - 1.69) |
| **rs2242549** | 0.56 | 0.74 | 1.10 (0.64 - 1.86) |
| **rs881803** | 0.29 | 0.15 | 1.53 (0.86 - 2.73) |
| **rs2075512** | 0.64 | 0.79 | 0.93 (0.53 - 1.63) |
| **Haplotype** | 0.35 | 0.73 | 1.10 (0.63 - 1.93) |
| SSRIs (N=357-402, no of instances: 496-563) |  |  |  |
| **rs4781678** | 0.31 | 0.033 | 0.43 (0.20 - 0.93) |
| **rs2242549** | 0.92 | 0.031 | 0.46 (0.23 - 0.93) |
| **rs881803** | 0.41 | 0.29 | 0.67 (0.32 - 1.40) |
| **rs2075512** | 0.97 | **0.010** | **0.37 (0.17 - 0.79)** |
| **Haplotype** | 0.87 | 0.029 | 0.42 (0.19 - 0.91) |
| SSRIs metabolised by CYP2C19 (N=282-318, no of instances: 357-395) |  |  |  |
| **rs4781678** | 0.42 | **0.003** | **0.27 (0.11 - 0.64)** |
| **rs2242549** | 0.47 | **0.006** | **0.30 (0.13 - 0.70)** |
| **rs881803** | 0.51 | 0.064 | 0.44 (0.19 - 1.05) |
| **rs2075512** | 0.39 | **0.005** | **0.31 (0.13 - 0.70)** |
| **Haplotype** | 0.95 | **0.006** | **0.30 (0.13 - 0.71)** |
| SSRIs not metabolised by CYP2C19 (N=123-145, no of instances: 142-169) |  |  |  |
| **rs4781678** | 0.48 | 0.71 | 1.36 (0.27 - 6.87) |
| **rs2242549** | 0.30 | 0.58 | 1.37 (0.45 - 4.23) |
| **rs881803** | 0.58 | 0.23 | 2.24 (0.60 - 8.38) |
| **rs2075512** | 0.10 | 0.88 | 0.88 (0.19 - 4.13) |
| **Haplotype** | 0.66 | 0.77 | 1.25 (0.27 - 5.76) |
| Non-SSRIs metabolised by CYP2C19 (N=318-355, no of instances: 411-459) |  |  |  |
| **rs4781678** | 0.24 | **0.0051** | **3.33 (1.44 - 7.73)** |
| **rs2242549** | 0.62 | **0.0013** | **3.42 (1.61 - 7.25)** |
| **rs881803** | 0.63 | **0.00030** | **5.82 (2.25 - 15.0)** |
| **rs2075512** | 0.92 | **0.0064** | **3.63 (1.44 - 9.17)** |
| **Haplotype** | 0.22 | **0.00030** | **4.93 (2.07 - 11.76)** |

### The Effect of miR-484 on CYP2C19 in Cultured Cells

Given that a major effect of the *NDE1* locus variants examined here seems to be to alter the expression of genes targeted by miR-484, presumably due to altered expression of this miR-484, we hypothesized that the pharmacological consequences of these variants were also likely to occur through miR-484. For this to be the case, miR-484 would need to affect the levels of CYP2C19 protein expression, and thus be able to alter its metabolic activity effects on psychoactive medication.

We therefore conducted a proof of principle experiment in NLF human neuroblastoma cells, into which we transfected a mimic of the mature form of human miR-484. Protein levels of CYP2C19 were significantly up-regulated following miR-484 transfection, when compared to transfection with a negative control microRNA (Figure 2). In contrast, no effect on the expression of another principal metabolizing enzyme of psychoactive medications, CYP2D6, was seen, indicating that this is a specific effect.

**Figure 2:**
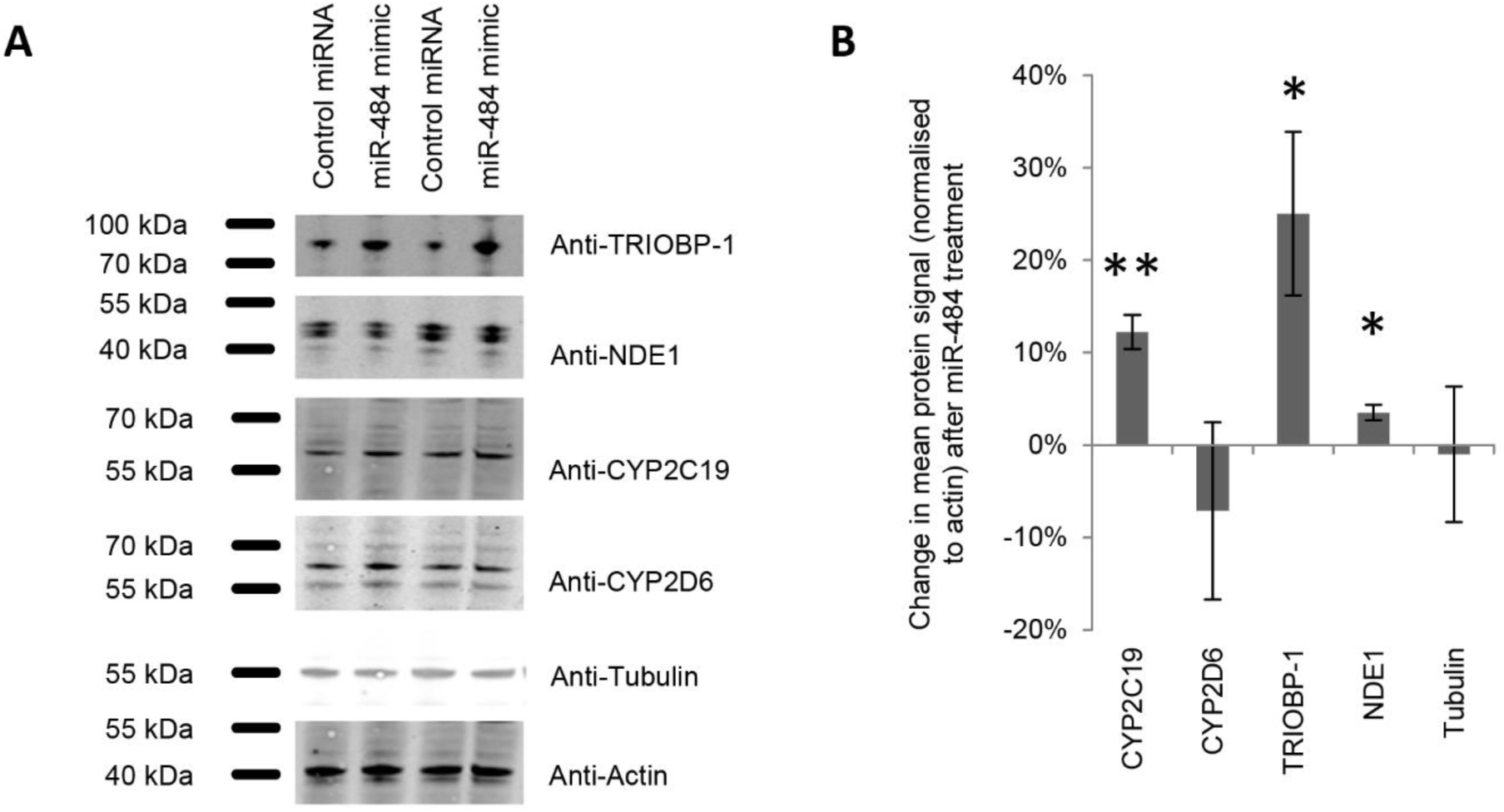
The effect of a miR-484 mimic on protein expression in human NLF neuroblastoma cells. a) Sample western blots showing levels of six proteins in the lysates of cells which had been transfected with either a mimic of miR-484 or with a negative control microRNA. b) Quantification of three independent experiments, each comprising 7-8 internal replicates. All proteins were normalized to actin. ^*^: p < 0.05, ^**^: p < 0.01.

Finally, we also used the same system to investigate two proteins, each of which illustrates a potential false negative in this study, possibly as a result of power limitations. The first of these, *TRIOBP* was observed to be significant in our discovery cohort but not in the replication cohorts, while the other is *NDE1* itself, which was highly significant in the larger GTEx database (beta = 0.24; Standard error = 0.037; t-statistic=6.6; p=2.3x10^−10^), but not originally observed in our discovery cohort. *TRIOBP* was further selected as it is also an example of the six genes whose expression was associated with *NDE1* variation both here and in our previous analysis (30). In both cases, proteins levels were subtly, but statistically significantly altered following treatment with the miR-484 mimic (Figure 2), in comparison to the negative control microRNA.

## Discussion

Here we have demonstrated that variations within the *NDE1* locus, encoding a protein of the DISC1 network of protein interaction partners, can affect both gene expression levels and medication usage of psychoactive drugs used to treat major mental illnesses. Specifically, two SNPs in high LD are associated with replicable expression changes in a large number of genes, and with early cessation of psychoactive medications metabolised by CYP2C19 in a gender dependent manner. We propose that these observations are linked through the involvement of miR-484. This microRNA is encoded for within the 5’ untranslated exon of the longest splice variant of *NDE1,* and the one which is most abundantly expressed, at least in cell culture (50). Notably, the list of genes whose expression changes are associated with these *NDE1* locus variants is significantly enriched for predicted targets of the microRNA, while expression of the CYP2C19 protein has been demonstrated *in vitro* to be significantly increased following addition of a mimic for mature human miR-484. The most promising explanation for the observations described here would therefore be that variation at the *NDE1* locus affects gene expression and medication metabolism in large part through effects of the variant on miR-484, and this may even be behind our prior observations at this locus of association to schizophrenia. It is interesting to note that a 1.45-fold increase in miR-484 has previously been reported in the superior temporal gyri of patients with schizophrenia (51).

The 16p13.11 locus, in which the *NDE1* gene is found, is prone to copy number variations (CNVs), with these 16p13.11 CNVs having been repeatedly associated with psychiatric and neurological disorders (19-22, 28). This locus contains multiple genes, however *NDE1* has been considered amongst the most promising candidates to be involved in these disorders due to its known critical role in neurodevelopment (reviewed: 52). Therefore, our observations here, although of specific SNPs at the *NDE1* locus, highlights disruption of miR-484 as a potential functional consequence also of those CNVs. These results partially parallel recent findings that expression of mature miR-484 led to alterations in neural progenitor proliferation and differentiation, as well as behavioural changes in mice, thus implicating the microRNA in the phenotypes associated with 16p13.11 duplication (29). While NDE1 over-expression was not seen to have a gross effect on neuronal progenitor proliferation under similar circumstances, given the severe neurological consequences of biallelic disruption of the *NDE1,* but not miR-484, reading frame (24-27), there is still a potential role for NDE1 in the conditions associated with 16p13.11 duplication. Additionally, relatively mild phenotypic effects would be needed to explain the fact that while associated with schizophrenia risk, most carriers of the CNV do not develop the condition (19-22). Nevertheless, it can be speculated that a consequence of the duplication of this locus would be gene expression levels changes driven by miR-484, as was seen here to be the case with *NDE1* SNP rs2242549.

This study initially sought to replicate our previous work on the effect of DISC1 network variants on gene expression changes in the general population, using publicly available data on the CEU (Utah residents with North and Western European ancestry) individuals (30). Of these previously identified 86 genes, we were able to replicate the observed changes in expression of six genes, including expression changes of two probes for *ST3GAL5* and a probe for *TRIOBP,* all of which were in association with the *NDE1* SNP rs2242549. When we tested for gene expression alterations across the genome we identified a large number of probes (n=2,908) representing 2,542 genes whose expression levels associated with variants at the *NDE1* locus, specifically with the SNP rs2242549. A large proportion of these (752 out of 3,824 probes, 695 out of 3,314 genes) were significantly altered by another *NDE1* SNP (rs1050162), which is in high LD with rs2242549. Yet replication attempts in independent cohorts, although providing validation for some genes (56 in an independent Finnish schizophrenia cohort and 180 using the GTEx database), did not provide unilateral confirmatory evidence, with the exception of 4 genes *(ITGB5, OVGP1, PGRMC1, TST)* identified across all three datasets tested. This lack of replication, combined with new observations and their biological relevance through miR-484 to the *NDE1* locus, suggests that, although the variants studied here are common to many populations, their relationship to potential functional mutations at this locus, and their specific biological consequences associated with schizophrenia and gene expression changes may be unique to this Finnish family cohort (53). This population difference may account for the lack of replication of most of the previously observed genes in the CEU population (30), and the lack of enrichment for miR-484 targets in the GTEx database. This is consistent with *DISC1* variation playing a genetically heterogeneous role in the general incidence of schizophrenia, lacking common illness-associated variations which could be detected by genome-wide association studies (15, 16) of global populations, but providing strong evidence for a role in the condition within specific populations and family studies (53).

Another potential explanation for our inability to replicate our observations across cohorts is the fact that our power to detect these effects is reduced due to the small sample sizes used here. Although we have demonstrated that we have 80% power in our discovery cohort to detect large changes in gene expression (Δ =0.52), this is for our observed 90^th^ percentile of the standard deviation for all genes from our data (σ= 0.53) (Figure S2), probes with smaller standard deviations would not be detectable, either for these probes in replication cohorts or for other probes in the discovery cohort. Thus, we verified our observations in a neuroblastoma cell culture model for two genes that provided inconsistent observations. The first, *TRIOBP,* was observed in our previously published study of the publicly available data on the CEU (Utah residents with North and Western European ancestry) individuals (30) and was replicated in our discovery family cohort, but not in either the twin or GTEx replication cohorts. In contrast, the second, *NDE1,* was not observed in any of the Finnish cohorts but was strongly implicated in the larger GTEx data where *NDE1* expression levels were strongly associated with the *NDE1* SNP rs2242549 genotype (beta = 0.24; Standard error = 0.037; t-statistic=6.6; p=2.3×10^−10^). The proteins encoded for by these genes were each found to be significantly increased by the presence of the mimic miR-484. Such a verification analysis would be required for all genes implicated in this study. However, with such a large number of genes identified this was not feasible with the cell culture model used here.

When the DISC1 network was studied with respect to treatment, the *NDE1* locus again demonstrated association, specifically in interaction with gender for drugs metabolized by CYP2C19. The degree of expression of cytochrome P450 enzymes in lymphocytes was too low to allow us to investigate potential changes in expression level in our family data. Instead we demonstrated in a cell culture model that miR-484 is capable of increasing the expression of CYP2C19, but not that of another major metabolizing enzyme for psychoactive drugs, CYP2D6. This suggests that the mechanism through which the locus confers risk and alters medication usage is the same. In the case of medication use, we employed a dichotomous variable based on a cut-off of ceasing to use the prescribed medication after three months or less. This was designed to indicate that a treatment was either not considered to be working or else was having side effects which were too severe and its use was therefore stopped. Since the cell culture experiment showed that CYP2C19 protein expression is increased by miR-484, it can be hypothesized that the medications are more rapidly metabolized in individuals carrying these variants, leading to a reduced efficacy of those treatments. Interestingly, the genetic effects on medication differ depending on both class of drug and gender. Males carrying the risk alleles had a higher probability of cessation of use for SSRIs metabolized by CYP2C19, while females carrying the risk alleles had an increased probability of cessation for non-SSRI drugs that are metabolized by CYP2C19. Although the mechanism for these effects remains unclear at this time, it is noteworthy that the original association between the *NDE1* locus and schizophrenia in these families was significant only in females (3). Taken together this implies that one or more gender-specific effects act as modifying factors in conjunction with the *NDE1*/miR-484 locus, although this cannot be easily modelled in our cell culture system.

Here, through the identification of altered gene expression patterns that led to the functional implication of miR-484, which is coded on an untranslated exon of *NDE1*, we identified a means by which genetic variation in the DISC1 network can not only increase risk to major mental illnesses, but also how those same variants can alter treatment response to specific psychoactive medications through the regulation of their metabolizing enzyme. This study has therefore provided new biological insight into psychiatric disorders to which novel medications could be designed, as well as suggesting that knowledge of an individual’s genotype within the *NDE1*/miR-484 locus may have potential value in the targeting of current therapies.

## Ethics

In all studies, the principles recommended in the Declaration of Helsinki, and its amendments were followed. The study has been approved by the Coordinating Ethics committee of the Hospital District of Helsinki and Uusimaa. Informed consent was obtained from all participants.

## Author Contributions

NJB, LUV and WH wrote the manuscript text; NJB, MP, LUV, and WH prepared the manuscript figures; NJB, JS, JL, JH and WH designed the study; ST, TP, JS, JL, TDC and JH provided access to samples and data; NJB performed laboratory experiments; NJB, LUV, MP, ABZ, AOA, MTH, VS and WH performed the analysis; all authors reviewed the manuscript and approved the final version to be published.

## Acknowledgements

Gene-expression analysis was performed by the Institute for Molecular Medicine Finland FIMM Technology Centre, University of Helsinki. Jaakko Kaprio is gratefully acknowledged for the provision of control twins to this study and for critical reading of the manuscript. Antti Tanskanen of the National Institute for Health and Welfare is thanked for producing the definitions of the medications periods used in this study.

